# Transcriptome analysis of FOXO-dependent hypoxia gene expression identifies Hipk as a regulator of low oxygen tolerance in *Drosophila*

**DOI:** 10.1101/2022.06.17.496640

**Authors:** Kate Ding, Elizabeth C Barretto, Michael Johnston, Byoungchun Lee, Marco Gallo, Savraj S Grewal

## Abstract

When exposed to low oxygen or hypoxia, animals must alter their metabolism and physiology to ensure proper cell-, tissue- and whole-body level adaptations to their hypoxic environment. These alterations often involve changes in gene expression. While extensive work has emphasized the importance of the HIF-1 alpha transcription factor on controlling hypoxia gene expression, less is known about other transcriptional mechanisms. We previously identified the transcription factor FOXO as a regulator of hypoxia tolerance in *Drosophila* larvae and adults. Here we use an RNA-sequencing approach to identify FOXO-dependent changes in gene expression that are associated with these tolerance effects. We found that hypoxia altered the expression of over 2000 genes and that approximately 40% of these gene expression changes required FOXO. We discovered that hypoxia exposure led to a FOXO-dependent increase in genes involved in cell signaling, such as kinases, GTPase regulators, and regulators of the Hippo/Yorkie pathway. Among these, we identified homeodomain-interacting protein kinase (Hipk) as being required for hypoxia survival. We also found that hypoxia suppresses the expression of genes involved in ribosome synthesis and egg production, and we showed that hypoxia suppresses tRNA synthesis and mRNA translation and reduces female fecundity. Among the downregulated genes, we discovered that FOXO was required for suppression of many ribosomal protein genes and genes involved in oxidative phosphorylation, pointing to a role for FOXO in limiting energetically costly processes such as protein synthesis and mitochondrial activity upon hypoxic stress. This work uncovers a widespread role for FOXO in mediating hypoxia changes in gene expression.

## Introduction

Animals often live in conditions where environmental oxygen levels fluctuate (Clegg 1997; Danovaro *et al*. 2010; Park *et al*. 2017). As a result, they must coordinate their physiology and metabolism with changes in oxygen availability to maintain proper homeostasis. This coordination can occur through alterations in gene expression and is essential for ensuring organismal survival in low oxygen (Bickler and Buck 2007; Ramirez *et al*. 2007; Harrison and Haddad 2011; Padilla and Ladage 2012; Nakazawa *et al*. 2016; Samanta *et al*. 2017; Harrison *et al*. 2018; Schito and Rey 2018; Holdsworth and Gibbs 2020).

Across all metazoans, perhaps the best-described and most intensively studied mechanism of gene regulation in hypoxia involves the HIF-1 alpha transcription factor (Semenza 2011; Semenza 2014b). When cells encounter low oxygen conditions, HIF-1 alpha protein is stabilized and translocates to the nucleus to induce gene expression. HIF-1 alpha-regulated genes encompass a diverse array of genes that are involved in biological processes such as metabolism, cell signaling and transcription, and that together coordinate cell-, tissue- and whole-body level adaptations to low oxygen (Semenza 2011; Samanta *et al*. 2017). Studies in model organisms have identified how HIF-1 alpha is a key regulator of hypoxia in both normal physiology and in pathological disease states (Semenza 2014a; Semenza 2014b). However, compared to our understanding of HIF-1 alpha biology, less is known about other transcriptional mechanisms that contribute to both cellular and systemic oxygen homeostasis.

*Drosophila* have provided a versatile and informative model system for investigating organismal responses to hypoxia. In their natural ecology, *Drosophila* live and grow on rotting, fermenting food rich in microorganisms – an environment characterized by low ambient oxygen (Callier *et al*. 2015; Markow 2015; Harrison *et al*. 2018). They have therefore evolved mechanisms to tolerate hypoxia. For example, larvae and adults can tolerate severe hypoxia (∼1% oxygen) for up to 24 hours with little impact on viability (Barretto *et al*. 2020), while embryos can survive complete anoxia (0% oxygen) for several days (Foe and Alberts 1985; Teodoro and O’Farrell 2003). Genetic studies have shown that flies can survive oxygen deprivation by increasing tracheal branching to expand oxygen supply to tissues (Centanin *et al*. 2008; Wong *et al*. 2014), and by remodelling their physiology and metabolism through both HIF-1 alpha-dependent and independent mechanisms (Wingrove and O’Farrell 1999; Lavista-Llanos *et al*. 2002; Teodoro and O’Farrell 2003; Centanin *et al*. 2005; Romero *et al*. 2007; Harrison and Haddad 2011; Morton 2011; Li *et al*. 2013; Bandarra *et al*. 2014; Harrison *et al*. 2018; Lee *et al*. 2019; Texada *et al*. 2019; Barretto *et al*. 2020). Relatively few studies, however, have used genome-wide approaches to identify gene expression changes associated with adaptation to hypoxia in *Drosophila* (Liu *et al*. 2006a; Li *et al*. 2013). One study examined transcriptome changes associated with larval hypoxia and identified widespread changes in metabolic and gene expression (Li *et al*. 2013). This study also showed that of the hundreds of gene expression changes, over half were independent of HIF-1 alpha, emphasizing the importance of additional transcriptional mechanisms in the control of hypoxia gene expression (Li *et al*. 2013).

Using *Drosophila* larvae and adults, we previously identified the transcription factor, Forkhead Box O (FOXO), as a regulator of hypoxia tolerance (Barretto *et al*. 2020). FOXO is a conserved regulator of stress responses and animal aging (Webb and Brunet 2014). Studies in *Drosophila* have shown it is induced by stressors such as starvation, oxidative stress, pathogens, and ionizing radiation (Junger *et al*. 2003; Dionne *et al*. 2006; Karpac *et al*. 2009; Karpac *et al*. 2011; Borch Jensen *et al*. 2017). Genetic studies have also shown that, in general, loss of *foxo* induces stress sensitivity and shortens lifespan whereas increased FOXO activity, particularly in tissues such as gut, muscle and fat body can promote stress resistance and extend lifespan (Giannakou *et al*. 2004; Hwangbo *et al*. 2004; Tettweiler *et al*. 2005; Kramer *et al*. 2008; Demontis and Perrimon 2010; Alic *et al*. 2014a; Alic *et al*. 2014b; Dobson *et al*. 2017). We showed that FOXO activity is rapidly induced in hypoxia and that it is needed for hypoxia survival in both larvae and adults. We also identified the immune Relish/NF Kappa B transcription factor as one target of FOXO important for its hypoxia tolerance effects. However, it is unclear what other genes FOXO may regulate in hypoxia. Previous studies have shown that under normal conditions FOXO can bind to thousands of genomic loci (Alic *et al*. 2011; Birnbaum *et al*. 2019) and can regulate the expression of hundreds genes in a tissue- and context-specific manner (Alic *et al*. 2011; Alic *et al*. 2014a; Alic *et al*. 2014b), raising the possibility that it may mediate broad effects on gene expression in hypoxia.

In this report we describe our transcriptome analysis of hypoxia-mediated gene expression changes upon hypoxia in adult flies. We show that FOXO is required for upregulation of genes involved in cell signaling and we identify the kinase Hipk as a regulator of hypoxia tolerance. We also see that FOXO suppresses expression of genes involved in protein synthesis and mitochondrial activity, suggesting it plays an important role in limiting energetically costly processes in low oxygen stress.

## Material and Methods

### *Drosophila* Stocks and Culturing

Flies were grown on medium containing 150 g agar, 1600 g cornmeal, 770 g Torula yeast, 675 g sucrose, 2340 g D-glucose, 240 ml acid mixture (propionic acid/phosphoric acid) per 34 L water. All stocks were maintained at either 18°C or RT. For adult hypoxia exposures, flies were raised from embryos to adults at 25°C and then, following eclosion, females were allowed to mate for 2 days before being separated from males and aged for another 5-6 days, at which time point hypoxia experiments were performed. For larval hypoxia exposures, hatched larvae were grown on food at 25°C until 96hrs after egg-laying, at which time point hypoxia experiments were performed. The following *Drosophila* strains were used: *w*^*1118*^, *foxo*^*D94*^*/TM6B, UAS-hipk* RNAi, *da-GS-Gal4 (daughterless-GeneSwitch)*. For UAS gene induction using the GeneSwitch system, adult flies were fed food supplemented with RU486 (200µM) for seven days. Control (non-induced flies) were maintained on normal food supplemented with ethanol (vehicle control for RU486).

### Hypoxia exposure and measurement of hypoxia survival

Vials of adult flies or larvae were placed into an airtight glass chamber into which a mix of 1% oxygen/99% nitrogen gas continually flowed. Flow rate was controlled using an Aalborg model P gas flow meter. Normoxic animals were maintained in vials in ambient air. For hypoxia survival experiments, mated female adults were placed in placed into hypoxia (1% oxygen) for 20 hours in groups of 15 flies per vial. Then, vials were removed from hypoxia and the flies were allowed to recover before the numbers of dead flies were counted.

### Total RNA isolation

Adult flies (5 per group), adult tissues (from 10 animals per group), or larvae (10 per group) were snap frozen on dry ice. Total RNA was then isolated using Trizol according to the manufacturer’s instructions (Invitrogen; 15596-018). Extracted RNA was then DNase treated (Ambion; 2238G) to be used for subsequent qPCR or mRNA-sequencing.

### mRNA-sequencing and RNA-seq analyses

Three to four independent biological replicates (5 flies per group) of normoxia- and hypoxia-exposed groups of *w*^*1118*^ and *foxo* mutants were prepared and analyzed. RNA-sequencing was conducted by the University of Calgary Centre for Health Genomics and Informatics. The RNA Integrity Number (RIN) was determined for each RNA sample. Samples with a RIN score higher than 8 were considered good quality, and Poly-A mRNA-seq libraries from such samples were prepared using the Ultra II Directional RNA Library kit (New England BioLabs) according to the manufacturer’s instructions. Libraries were then quantified using the Kapa qPCR Library Quantitation kit (Roche) according to the manufacturer’s directions. Finally, RNA libraries were sequenced for 100 cycles using the NextSeq 500 Sequencing System (Illumina). Transcripts were quantified using kallisto (Bray *et al*. 2016) referencing refSeq mRNA (release: 2019-Oct-15) corresponding to dm6 annotation. Differential expression testing was performed using sleuth (Pimentel *et al*. 2017).

### Gene Ontology, KEGG pathway, and tissue expression analyses

Analyses of Gene Ontology and KEGG pathway enrichment of up- and down-regulated genes (>1.5-fold, q-val (FDR corrected p-val) <0.05) were performed using G-profiler (Raudvere *et al*. 2019) and Revigo (Supek *et al*. 2011).

### Quantitative RT-PCR measurements

Total RNA was extracted from either whole flies, whole larvae, or isolated adult tissues. The RNA was then DNase treated as described above and reverse transcribed using Superscript II (Invitrogen; 100004925). The generated cDNA was used as a template to perform qRT–PCRs (ABI 7500 real time PCR system using SyBr Green PCR mix) using gene-specific primers. PCR data were normalized to *beta tubulin* or *eIF2 alpha* mRNA levels. The following primers were used:

*tRNA ala* Forward: GCGGCCGCACTTTCACTGACCGGAAACG

*tRNA ala* Reverse: GCGGCCGCGCCCGTTCTAACTTTTTGGA

*tRNA arg* Forward: GCGGCCGCGTCCGTCCACCAATGAA AAT

*tRNA arg* Reverse: GCGGCCGCCGGCTAGCTCAGTCGGT AGA

*tRNA eMet* Forward: GCGGCCGCCGTGGCAATCTTCTGAA ACC

*tRNA eMet* Reverse: GCGGCCGCTCAGTGGAAAACCATA TGTTCG

*tRNA iMet* Forward: AGAGTGGCGCAGTGGAAG

*tRNA iMet* Reverse: AGAGCAAGGTTTCGATCCTC

*beta tubulin* Forward: ATCATCACACACGGACAGG

*beta tubulin* Reverse: GAGCTGGATGATGGGGAGTA

*Hipk* Forward: CAACAATGTCAAGGCATC

*Hipk* Reverse: CAGGCTGCACAGTGTGGAAA

*eIF2 alpha* Forward: TCTTCGATGAGTGCAACCTG

*eIF2 alpha* Reverse: CCTCGTAACCGTAGCAGGAG

### Statistical analysis of qRT-PCR data

All qRT-PCR data were analyzed by Student’s t-test or two-way ANOVA followed by post-hoc Students’ t-test where appropriate. All statistical analyses and data plots were performed using Prism statistical software. Differences were considered significant when p values were less than 0.05.

### Polysome Profiling

Larvae were lysed in lysis buffer (25 mM Tris pH 7.4, 10 mM MgCl_2_, 250 mM NaCl, 1% Triton X-100, 0.5% sodium deoxycholate, 0.5 mM DTT, 100 mg/ml cycloheximide, 1 mg/ml heparin, 16 Complete mini roche protease inhibitor, 2.5 mM PMSF, 5 mM sodium fluoride, 1 mM sodium orthovanadate and 200 U/ml ribolock RNAse inhibitor (Fermentas)) using a Dounce homogenizer. Lysates were then centrifuged at 15,000 rpm for 20 minutes and the supernatant was removed carefully using a fine syringe to avoid the floating fat content. For each condition, lysates containing 300 mg of total RNA were then layered on top of a 15–45% w/w sucrose gradient (made using 25 mM Tris pH 7.4, 10 mM MgCl_2_, 250 mM NaCl, 1 mg/ml heparin, 100 mg/ml cycloheximide in 12 ml polyallomer tube) and centrifuged at 37,000 rpm for 150 minutes in a Beckmann Coulter Optima L-90K ultracentrifuge using a SW-41 rotor. Polysome profiles were obtained by pushing the gradient using 70% w/v sucrose pumped at 1.5 ml/min into a continuous OD254 nm reader (ISCO UA6 UV detector).

### Fecundity Assay

1-2 day old virgin *w*^*1118*^ females were allowed to mate with males for 2 days and then the females were separated and either maintained in normoxia (controls) or exposed to hypoxia for either 8 or 12 hours before being returned to normoxia. Twenty four hours later, the females were transferred in groups of 3 to new vials and allowed to lay eggs for a 24 hour period (day 1), and then transferred to a second set of vials to lay eggs for a further 24 hour period (day 2). Fecundity was then assessed by measuring the number of viable pupae per female that emerged from eggs laid on day 1 and day 2.

## Results

### Hypoxia leads to upregulation of transcription factor and kinase gene expression

Adult mated *w*^*1118*^ (control) or *foxo*^*Δ94*^ (foxo null mutant)(Slack *et al*. 2011) females were either maintained in normoxia or exposed to hypoxia (1% oxygen) for 16 hours and we then isolated whole-body RNA for RNA-seq analysis (Figure 1A). We first examined the gene expression changes induced by hypoxia in the control animals. Using a cut-off of +/-1.5 fold and a false-discovery rate corrected p-value <0.05, we identified 1081 genes with reduced mRNA expression and 1257 genes with increased mRNA expression in *w*^*1118*^ animals (Figure 1B, Supplemental Table 1). Among the upregulated genes, we saw increased expression of several genes previously shown to be induced upon hypoxia exposure in larvae and/or adults. For example, we saw increased expression of the fly fibroblast growth factor homolog, *branchless (bnl)*, the glycolytic enzyme, *Lactate dehydrogenease (Ldh/ImpL3)*, and the transcriptional repressor, *hairy (h)*, each of which has been shown to be upregulated upon hypoxia exposure (Centanin *et al*. 2008; Zhou *et al*. 2008; Li *et al*. 2013) (Figure 1C). We previously showed hypoxia induces rapid nuclear localization and increased transcriptional activity of the transcription factor FOXO, which we found promoted hypoxia tolerance by increasing expression of the innate immune transcription factor Relish (*Rel*) (Barretto *et al*. 2020). Consistent with this, our transcriptome data showed that hypoxia led to increased expression of two FOXO target genes, *Thor* and *InR*, and increased expression of *Rel* and antimicrobial peptide genes (e.g., *CecA1, CecA2, AttB, AttA, Dro)*, which are known targets of Relish (Figure 1C). Together these changes in gene expression confirm that our low oxygen exposure protocol induced a robust hypoxic response. Two previous studies used genome-wide transcriptome analyses to examine hypoxia regulated genes in *Drosophila* (Liu *et al*. 2006a; Li *et al*. 2013). Like us, Liu *et al*. examined hypoxia in adult female flies, and they used DNA microarray hybridization to identify genes that showed significantly increased expression after 6 hours of severe (0.5% oxygen) hypoxia exposure. They identified 79 genes, of which 47 (59%) were also identified in our RNAseq analysis (significant overlap, p = 7.3 × 10^−29^) (Supplemental Table 1). Li *et al*. also used DNA microarray hybridization to detect hypoxia-regulated genes, in this case, in late L3 larvae using milder hypoxia (4% oxygen). They identified 627 significantly (p<0.01) upregulated (>1.5 fold) genes, of which 130 (21%) were also identified as upregulated in our RNA-seq analysis (significant overlap, p = 2.2 × 10^−19^), and they identified 417 significantly (p<0.01) downregulated (>1.5 fold) genes, of which 80 (19%) were also identified in our RNA-seq analysis (significant overlap, p = 4.8 × 10^−13^) (Supplemental Table 1). Thus, even though this study analyzed hypoxia at a different stage of the life cycle and at a different concentration of oxygen, one-fifth of the genes that were identified as being hypoxia-regulated in larvae were also identified in our study in adults. The differences in the number of genes identified in our study versus the previous studies likely reflect differences in biology between larvae and adults, and the greater sensitivity of RNA-seq approaches to detect differentially expressed genes compared to DNA microarrays.

**Figure 1.**
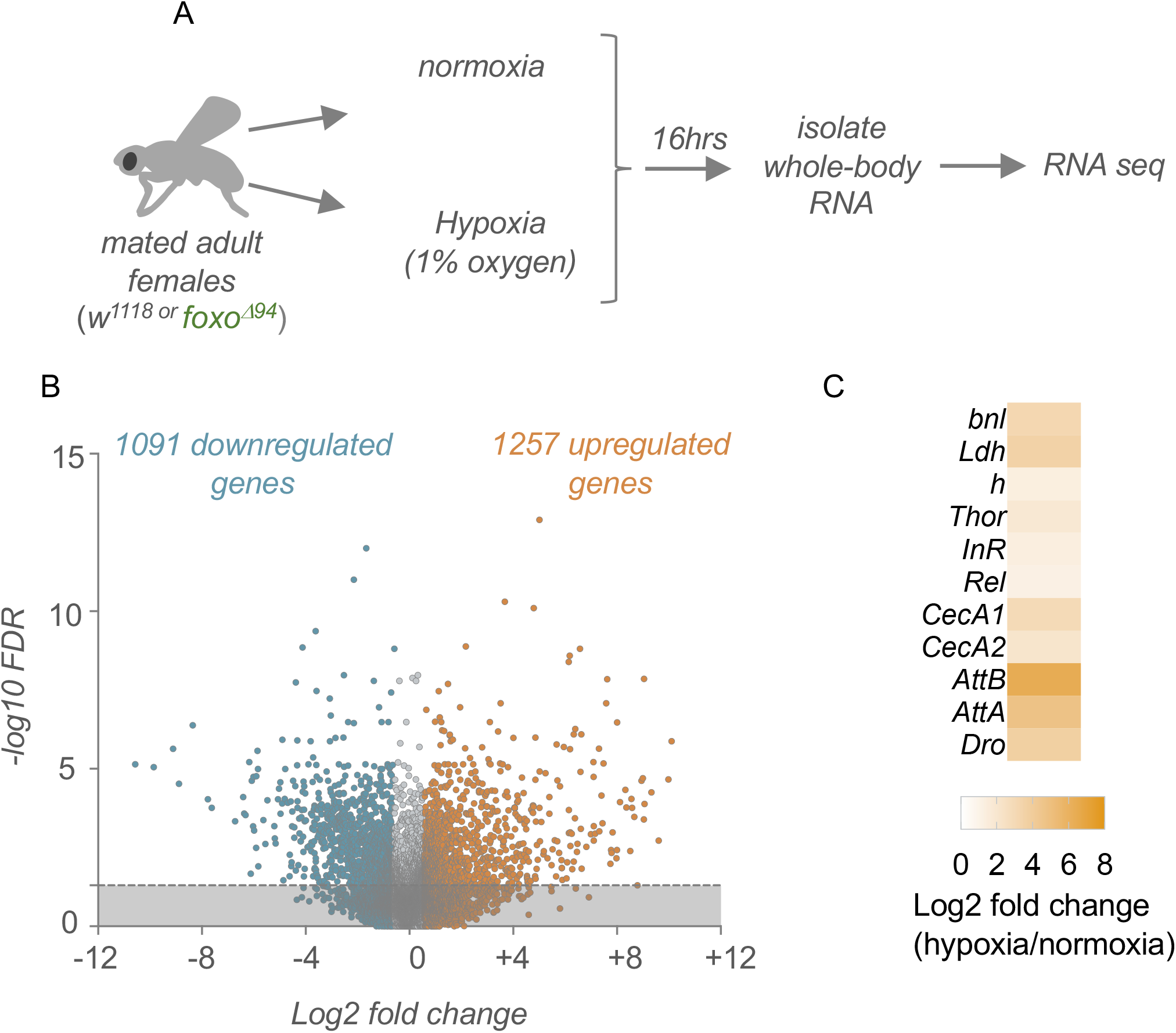
Hypoxia-induced alterations in whole-body gene expression. A) Schematic outline of our experimental approach. B) Volcano plot showing the up- (orange) and down-regulated (blue) genes following hypoxia exposure. Genes were considered differentially expressed if they showed a significant (q-val (FDR corrected p-val) <0.05) change in expression that was > +/- 1.5-fold different in hypoxia vs normoxia. Dashed line indicates q-val = 0.05. C) Heatmap depicting the change in expression (Log2 fold change, hypoxia vs normoxia conditions) of previously described hypoxia-induced genes.

We used Gene Ontology analysis to examine the genes that showed upregulated expression upon hypoxia. This analysis showed that the upregulated genes were particularly enriched for gene categories related to chromatin modification and transcription, small G-protein regulators, and kinases (Figure 2A). In addition, KEGG pathway analysis of the upregulated genes showed enrichment for genes involved in Hippo, Notch, FOXO, and MAPK signaling (Figure 2B). We saw hypoxia-induced increases in gene expression for 132 regulators of transcription and chromatin, and 61 kinases (Figure 2C). Together, these analyses suggest that hypoxia leads to widespread upregulation of different signaling pathways and transcriptional responses.

**Figure 2.**
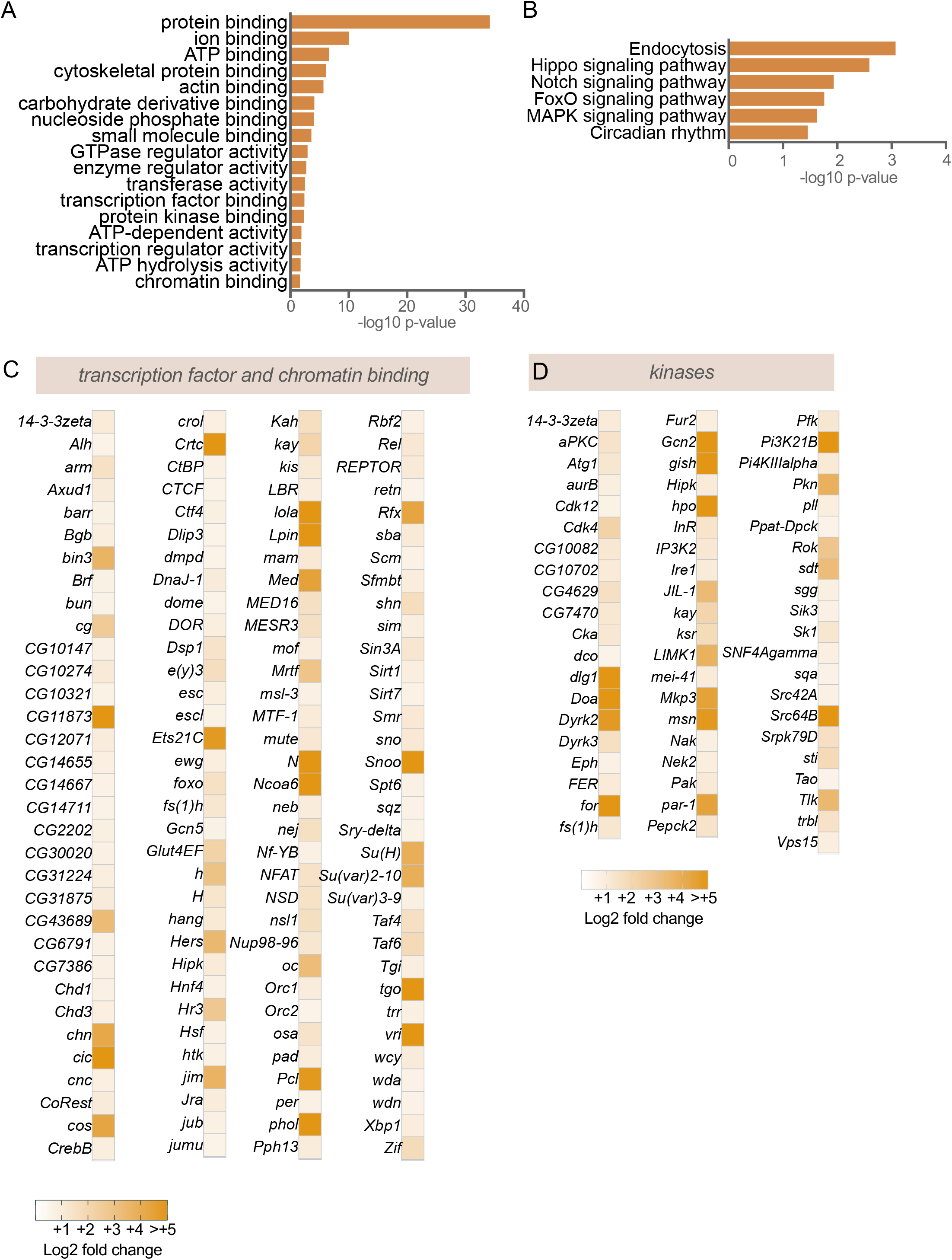
Hypoxia upregulates mRNA expression of transcription factors and kinase genes. A) GO analysis (molecular function category), and B) KEGG pathway analysis of genes showing >1.5-fold increase following hypoxia exposure. C) Heatmap depicting the increases in mRNA expression (Log2 fold change hypoxia vs normoxia) of transcription factor genes. D) Heatmap depicting the increases in mRNA expression (Log2 fold change hypoxia vs normoxia) of kinase genes.

### FOXO is required for hypoxia-induced upregulation of signaling molecules and regulators of the Hippo pathway

To identify FOXO-dependent hypoxia-induced genes we identified genes that showed a significant (q-val <0.05), >1.5-fold upregulation upon hypoxia in *w*^*1118*^ but not *foxo*^*Δ94*^. Using this criteria, we found that of the 1257 genes that were upregulated upon hypoxia exposure, 551 (44%) were not significantly upregulated in *foxo* mutants, suggesting that a large proportion of hypoxia-induced gene expression requires FOXO activity (Supplemental Table 1). Two previous studies used ChIP-chip and ChIP-seq approaches to identify FOXO genomic binding sites in young female adult flies, and, between them, identified 3925 loci that bound FOXO and were within 1kb of a protein coding gene (Alic *et al*. 2011; Birnbaum *et al*. 2019). Interestingly, we saw that of the 553 FOXO-dependent hypoxia upregulated genes that we identified, 265 (48%) overlapped with these FOXO-bound genes (Figure 3A), suggesting that almost half the FOXO-dependent hypoxia genes may be induced by direct FOXO transcriptional activation (Supplemental Table 1). We used Gene Ontology analysis to examine the 553 FOXO-dependent hypoxia-upregulated genes. The main classes of genes identified were largely related to signaling regulators, such as kinases, GTPase regulators, and guanine-nucleotide exchange factors (Figure 3C). For example, we saw that many GTPase regulators required FOXO for their hypoxia upregulation and many of these contained FOXO binding sites within 1Kb of their gene coding region. In addition, almost half the kinases that we saw were induced in hypoxia were dependent on FOXO for their induction. Interestingly, among these signaling molecules, we saw enrichment in regulators of the Hippo signaling pathway, many of which were previously shown to be enriched among FOXO-bound genes, suggesting that they may be directly regulated by FOXO (Figure 3D).

**Figure 3.**
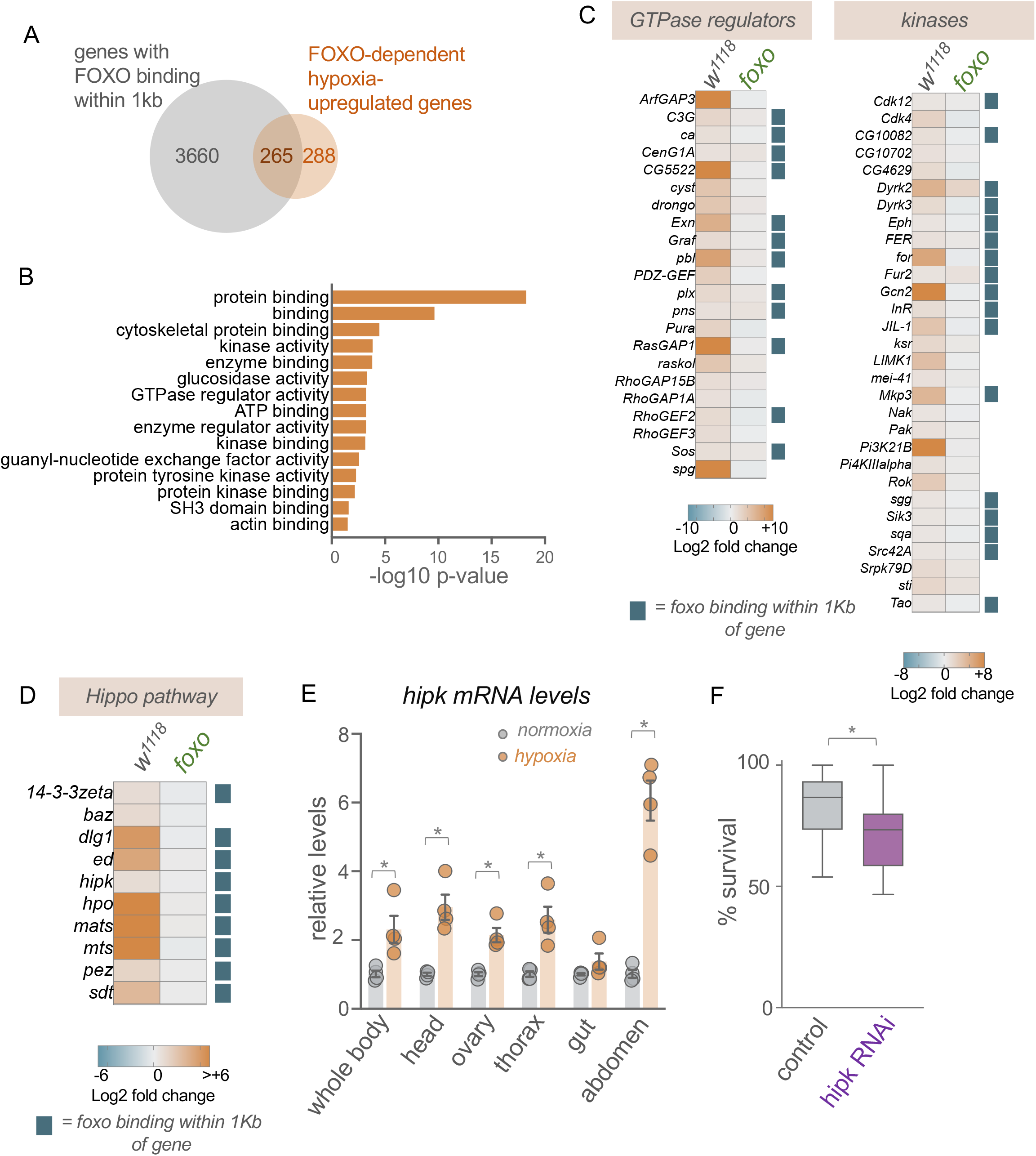
Hipk is a hypoxia-induced gene required for organismal hypoxia tolerance. A) Venn diagram showing overlap between genes previously shown to have FOXO binding within 1kb, as detected by ChIP, and FOXO-dependent upregulated genes identified in the present study. B) GO analysis (molecular function category) of FOXO-dependent hypoxia induced genes (genes showing a significant >1.5-fold increase in mRNA expression following hypoxia exposure in *w*^*1118*^ but not *foxo* mutants). C) Heatmap depicting the increases in mRNA expression (Log2 fold change, hypoxia vs normoxia) of GTPase regulators and kinases in *w*^*1118*^ and *foxo* mutants. Blue squares indicate genes previously shown to have FOXO binding within 1kb of the gene as measured by ChIP. D) Heatmap depicting the increases in mRNA expression (Log2 fold change, hypoxia vs normoxia) of Hippo pathway genes in *w*^*1118*^ and *foxo* mutants. Blue squares indicate genes previously shown to have FOXO binding within 1kb of the gene as measured by ChIP. E) qPCR analysis of *hipk* mRNA levels from normoxia vs hypoxia exposed animals. RNA was isolated from either whole animals or specific tissues. Bars represent mean +/- SEM. Symbols represent individual data points, n=4 per condition. * p<0.05, Students t-test. F) Hypoxia survival of control (*daGSG > hipk RNAi*, no RU486) vs *hipk* RNAi (*daGSG > Hipk RNAi*, RU486-treated) adult flies. Data are presented as box plots (25%, median and 75% values) with error bars indicating the min and max values, n = 14 groups of flies per condition.

### Hipk is upregulated in hypoxia and modulates hypoxia tolerance

One regulator of the Hippo pathway we saw upregulated and associated with a FOXO DNA binding site was Homeodomain interacting protein kinase (Hipk). In *Drosophila*, Hipk has been shown to control metabolism and growth in epithelial tissues, and has been shown to function as a regulator of several signaling pathways including Hippo, Wingless, Notch, JAK/STAT and JNK (Lee *et al*. 2009a; Lee *et al*. 2009b; Chen and Verheyen 2012; Poon *et al*. 2012; Verheyen *et al*. 2012; Blaquiere *et al*. 2014; Tettweiler *et al*. 2019; Wong *et al*. 2019; Kinsey *et al*. 2021; Steinmetz *et al*. 2021). In addition, a recent study in *C. elegans* showed that the worm homolog of Hipk, hpk1, was a regulator of worm survival in low oxygen (Doering *et al*. 2022). We therefore examined the role of Hipk in *Drosophila* hypoxia in more detail. Using qRT-PCR we confirmed that hypoxia exposure led to an increase in *hipk* mRNA levels in whole animals. We also saw hypoxia-induced increases in *hipk* mRNA levels in specific tissues such as the head, thorax (which is enriched in muscle), ovaries, and abdomen (which is enriched in adipose tissues), suggesting that the hypoxia-mediated increase in Hipk expression occurred across many tissues (Figure 3E). To explore the functional role for Hipk in hypoxia, we used the RNAi to knockdown *hipk* in flies and examined the effects on hypoxia tolerance. We used the *daughterless-GeneSwitch-Gal4 (da-GSG)* driver to induce ubiquitous expression of the dsRNA and to restrict RNAi-mediated knockdown of *hipk* to adult stages. We fed *da-GSG>hipk RNAi* females either normal food (control) or RU486-containing food to induce RNAi (*hipk RNAi*) and then examined the effects on hypoxia survival. We found that following 20 hours of hypoxia exposure the *hipk RNAi* animals had significantly reduced hypoxia survival compared to the control flies, suggesting that Hipk is required for hypoxia tolerance (Figure 3F).

### Hypoxia downregulates expression of protein synthesis and egg production genes and leads to reduced translation and fecundity

We used Gene Ontology analysis to examine the genes that showed significantly reduced (>1.5 fold decrease) expression upon hypoxia in control *w*^*1118*^ animals. We saw enrichment in genes involved in ribosome function, egg formation, and proteolysis (Figure 4A, B). Almost all the proteolysis genes were proteases that showed enriched expression in either the intestine or fat body (Supplemental Table 1). The decreased expression of genes involved in ribosome function is consistent with suppressed protein synthesis, a widely seen response to hypoxia in different organisms. We previously showed that regulation of tRNA synthesis was a key mechanism for regulating protein synthesis in *Drosophila*, particularly in response to nutrient starvation (Marshall *et al*. 2012; Rideout *et al*. 2012; Sriskanthadevan-Pirahas *et al*. 2018). When we examined tRNA levels by qPCR, we saw a strong reduction following hypoxia exposure (1% oxygen) in adults (Figure 4C). Furthermore, we saw that exposure of larvae to 1% oxygen also led to a strong reduction in tRNA levels that was observed at both 2 and 24 hours of hypoxia exposure (Figure 4D, E). We also saw that hypoxia larvae showed a similarly rapid decrease in overall translation compared to normoxic animals as shown by a decrease in polysome:monosome ratios in polysome profiles from whole animal lysates (Figure 4F). These results indicate that global suppression of protein synthesis is a common response to extreme hypoxia in both larvae and adults.

**Figure 4.**
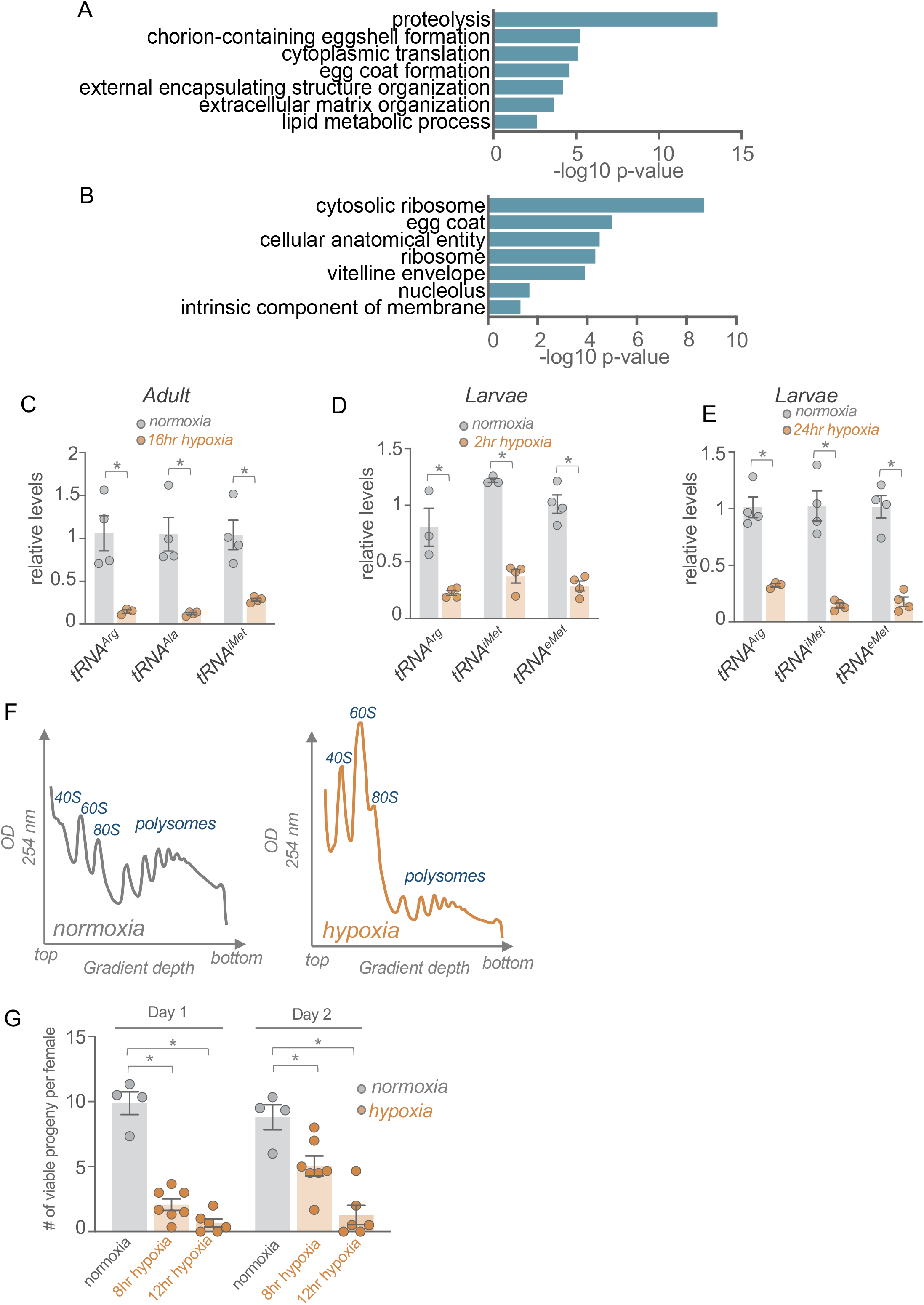
Hypoxia downregulates mRNA expression of protein synthesis and egg formation genes and leads to reduced translation and decreased fecundity. A, B) GO analysis (A, biological process category and B, cellular component category) of genes showing >1.5-fold decrease in expression following hypoxia exposure. C, D) qRT-PCR measurement of tRNA levels following C, 2hrs or D, 24hrs of hypoxia exposure in developing larvae. Bars represent mean +/- SEM. Symbols represent individual data points, n=4 per condition. * p<0.05, Students t-test. F) Polysome profiles of normoxia (left) and hypoxia (right) exposed larvae. Plots indicate continuous OD 254nm measurements from fractionated whole-body lysates. Peaks corresponding to 40S, 60S, 80S and polysomes are indicated. The top and bottom lysate fractions from the centrifuged sucrose gradients are indicated. G) Fecundity measurements from mated females exposed to normoxia or 8 or 12 hrs of hypoxia. Data show the mean number of viable pupae per female that developed from eggs laid on day 1 or day 2 following the hypoxia exposure. Bars represent mean +/- SEM. Symbols represent individual data points, n=4 per condition. * p<0.05, Students t-test following two-way ANOVA.

Given the decreased expression of egg formation genes, we also examined whether hypoxia might impact female fecundity. To do this, we exposed mated *w*^*1118*^ females to hypoxia for either 8 or 12 hours, allowed them to recover for a day, and then measured how many viable progeny they produced in the subsequent two days. We saw that when exposed to either 8 or 12 hours of hypoxia, females produced significantly fewer viable progeny compared to normoxic control females (Figure 4G). These results indicate that a brief exposure to hypoxia can suppress fecundity in female flies.

### FOXO is needed for hypoxia-mediated decreases in ribosome and mitochondrial gene expression

We then examined which downregulated genes were dependent on FOXO, by identifying genes that were significantly downregulated (<1.3 fold) in *w*^*1118*^ flies but not *foxo* mutants. We chose a lower fold change value because the significantly downregulated genes tended to be less affected than the upregulated genes. From this analysis, we identified 529 genes (39% of 1343 total downregulated genes) (Supplemental Table 1). Of these, 87 were previously shown to be bound to FOXO (Alic *et al*. 2011; Birnbaum *et al*. 2019), suggesting that FOXO-mediated decreases in gene expression in hypoxia are largely indirect (Figure 5A) (Supplemental Table 1). We used GO analysis to examine the functional categories of FOXO-dependent downregulated genes and identified strong enrichment in two main classes – ribosomal proteins and mitochondrial regulators (Figure 5B). We saw that genes coding for ribosomal proteins for both the small and large subunits showed reduced expression in *w*^*1118*^ but not *foxo* mutant animals (Figure 5C). We also found that many mitochondrial genes required FOXO for their downregulation in hypoxia, including known or predicted mitochondrial ribosome proteins, cytochrome-C oxidase subunits, complex I subunits, ATP synthases subunits, and mitochondrial transporters (Figure 5C). These results suggest that an important role for FOXO in hypoxia is to suppress both mitochondrial and protein synthetic activity.

**Figure 5.**
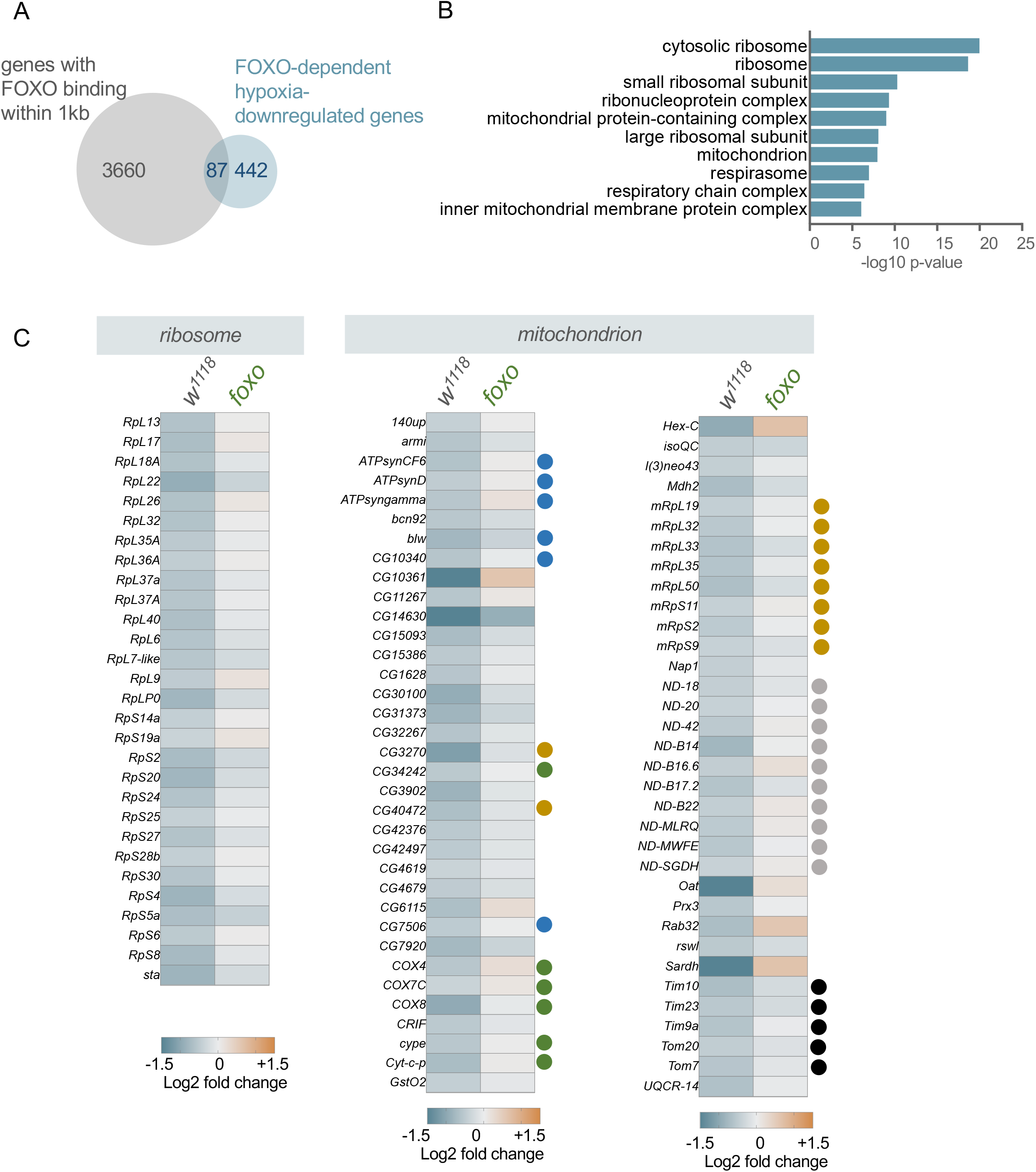
Hypoxia downregulation of ribosomal protein and mitochondrial regulator gene expression requires FOXO. A) GO analysis (cell component category) of FOXO-dependent hypoxia suppressed genes (genes showing a significant decrease in mRNA expression following hypoxia exposure in *w*^*1118*^ but not *foxo* mutants). B) Heatmap depicting the decreases in mRNA expression (Log2 fold change, hypoxia vs normoxia) of ribosomal protein genes and mitochondrial regulator genes in *w*^*1118*^ and *foxo* mutants. Colored circles indicate different classes of mitochondrial genes (blue: ATP synthase subunits; orange: mitochondrial ribosomal proteins; green: Cytochrome C oxidase subunits; grey: Complex I subunits; black: mitochondrial transporters).

## Discussion

We previously showed that the FOXO transcription factor was required for hypoxia tolerance (Barretto *et al*. 2020). One focus of this current study was to identify which genes might be FOXO-regulated in hypoxia. Our results indicate that hypoxia exposure alters (+/- 1.5 fold or greater) the transcript levels of ∼2300 genes in our control (*w*^*1118*^) line, indicating a widespread modification of gene expression. To identify which gene expression changes are FOXO dependent, we chose to identify which genes had significantly altered expression in *w1118* but not *foxo* mutants. This analysis showed that approximately 40% of hypoxia regulated genes required FOXO. Furthermore, using data from previous genome wide FOXO ChIP studies (Alic *et al*. 2011; Birnbaum *et al*. 2019) we saw that approximately half the FOXO dependent upregulated genes were directly bound by FOXO. These results suggest that FOXO is needed for widespread transcriptional changes upon hypoxia. A previous report examining genome-wide changes in gene expression upon hypoxia exposure in larvae showed that HIF-1 alpha was required for just under half of the changes in gene expression, and that the transcription factor estrogen-related receptor (ERR) was also important for mediating many of the effects of hypoxia on gene expression (Li *et al*. 2013). This study and our findings suggest that HIF-1 alpha, ERR, and FOXO may mediate many of the widespread changes in gene expression when flies are in low oxygen conditions. For example, we previously showed that FOXO was required for hypoxia tolerance in larvae (Barretto *et al*. 2020), suggesting it may cooperate or work in parallel with HIF-1 alpha and/or ERR to regulate hypoxia-mediated changes in gene expression at this developmental stage. Interestingly, we also saw that ERR mRNA levels were significantly increased upon hypoxia in adults (1.48-fold), although this was below our cut-off of 1.5-fold. Nevertheless, this suggests that ERR may also be important for hypoxia-mediated gene expression changes in the adult.

Hypoxia-upregulated genes were enriched for kinases, regulators of small GTPases, and regulators of gene expression such as transcription factors and chromatin modifiers. This suggests that a major response to hypoxia is widespread alterations in cell-cell signaling pathways and their downstream transcriptional effectors. We found that upregulation of many of these signaling genes was dependent on FOXO and likely direct, since many of these bound FOXO. Interestingly, regulators of the Hippo pathway were among the FOXO dependent upregulated genes. The Hippo pathway has been best studied in the context of cell growth and proliferation especially in epithelial, neural, and stem cells (Ma *et al*. 2019; Wu and Guan 2021). In these cells, the pathway often functions to couple cell-to-cell adhesion and cell polarity cues to the regulation of the downstream transcription factor Yorkie. Among the hypoxia-upregulated genes were several cell polarity/cell adhesion factors (*Ed, dlg1, sdt, baz)* and signaling molecules (*hpo, mats, pez, mts)* that function to negatively regulate Yorkie, suggesting that this may be an important regulator of hypoxia-mediated transcriptional responses. This regulation of Yorkie-mediated transcription may be important for regulation of stem or germ cell division upon hypoxia in adult flies. Yorkie can also regulate the processes of tracheal formation and immune signaling, which are both important in hypoxia. Recent studies have also shown that the mammalian homolog of Yorkie, Yap1, controls hypoxia-mediated angiogenesis in bone, suggesting that regulation of Hippo/Yorkie signaling may be a conserved hypoxia response (Sivaraj *et al*. 2020).

One kinase that showed FOXO-dependent increase in hypoxia was Hipk. We saw that this increase occurred across multiple tissues and was required for flies to survive hypoxia. These results point to Hipk as a regulator of hypoxia tolerance. As well as regulating Hippo/Yorkie signaling, Hipk can modulate other signaling pathways such as JNK, JAK/STAT, Wingless signaling(Lee *et al*. 2009b; Huang *et al*. 2011; Chen and Verheyen 2012; Poon *et al*. 2012; Verheyen *et al*. 2012; Blaquiere *et al*. 2014; Tettweiler *et al*. 2019; Kinsey *et al*. 2021; Steinmetz *et al*. 2021), as well as Notch signaling (Lee *et al*. 2009a), a pathway that we saw enriched in the KEGG analysis of hypoxia-upregulated genes. Thus, Hipk’s role in hypoxia tolerance may rely on regulation of any one of these pathways. Hipk has also been shown to induce glycolysis in larval epithelial tissues where it promotes tumor-like overgrowth (Wong *et al*. 2019). Hence, the hypoxia-mediated induction of Hipk may also be needed to induce glycolysis, a widely described metabolic response to low oxygen. Interestingly, a recent report showed that the *C. elegans* homolog of Hipk, hpk1, was needed for survival in low oxygen (Doering *et al*. 2022), suggesting a common role for Hipk in organismal hypoxia tolerance in both worms and flies.

Among the genes showing reduced expression in hypoxia, we saw strong enrichments for genes involved in egg production and translation. Furthermore, we saw that hypoxia suppressed female fecundity, and reduced translation and tRNA synthesis in both larvae and adults. Egg production is an energetically costly process, and therefore may be suppressed to ensure appropriate allocation of energetic resources to promote survival during stress. This type of trade-off between fecundity and stress responses has been seen in *Drosophila* in response to other environmental challenges. For example, upon infection with bacteria, fungi or viruses, flies have been shown to reduce their reproductive output and capacity (Schwenke *et al*. 2016). Moreover, germline deficient females that cannot produce eggs have enhanced immunity compared to fertile flies (Short *et al*. 2012). Similarly, nutrient starvation leads to reduced germline stem cell division and reduced egg production in females (Drummond-Barbosa and Spradling 2001; LaFever *et al*. 2010; Ables *et al*. 2012).

Protein synthesis is also an energetically costly process that has been estimated to account for at least one-third of a cell’s ATP use (Buttgereit and Brand 1995). Hence, it is not surprising that suppression of protein synthesis is a conserved response to hypoxia that is seen in many animals and that can promote hypoxia tolerance (Hofmann and Hand 1994; Hochachka *et al*. 1996; Liu *et al*. 2006b; van den Beucken *et al*. 2006; Anderson *et al*. 2009; Scott *et al*. 2013). Our transcriptomic analyses suggest that one way that hypoxia suppresses protein synthesis is by reducing expression of ribosome protein genes via FOXO. We also saw that FOXO was required for hypoxia-mediated suppression of many mitochondrial genes, including mitochondrial ribosomal proteins, mitochondrial transporters, and regulators of oxidative phosphorylation, such as subunits of ATP synthase, Cytochrome oxidase C, and Complex I. These results suggest that FOXO may also contribute to hypoxia tolerance by limiting energetically costly metabolic processes. FOXO suppression of ribosome protein and mitochondrial genes has also been seen in muscle following nutrient starvation in *Drosophila* larvae (Teleman *et al*. 2008). Furthermore, a recent study showed that the FOXO homolog in *C. elegans*, daf-16, promotes a hypoxia tolerant phenotype by suppressing ribosomal protein gene expression and partially suppressing genes involved in oxidative phosphorylation (Hemphill *et al*. 2022). Hence, reducing both ribosome gene expression and mitochondrial oxidative phosphorylation may be common FOXO-mediated stress responses.

In conclusion, our transcriptome analysis supports a model in which FOXO promotes hypoxia tolerance through controlling the upregulation of cell signaling pathways while suppressing the energetically costly processes of protein synthesis and mitochondrial activity. Given the conserved roles for FOXO in mediating hypoxia tolerance in different animals (Scott *et al*. 2002; Mendenhall *et al*. 2006; Menuz *et al*. 2009; Liu *et al*. 2016; Barretto *et al*. 2020; Hemphill *et al*. 2022) and the alterations of FOXO transcription factor activity in diseases associated with hypoxia, such as cancer, stroke, and ischemia (Maiese *et al*. 2008; Fukunaga and Shioda 2009; Maiese *et al*. 2009; Liu *et al*. 2022), our findings highlight processes that may contribute to low oxygen adaptations in both normal and disease states.

## Data availability

The RNA-sequence data has been deposited in NCBI’s Gene Expression Omnibus and are accessible through GEO Series accession number: GSE206206 (https://www.ncbi.nlm.nih.gov/geo/query/acc.cgi?acc=GSE206206)

## Acknowledgements

Stocks obtained from the Bloomington *Drosophila* Stock Center (NIH P40OD018537) were used in this study.

## Author contributions

K.D., E.C.B and B.L, carried out genetic and molecular experiments in *Drosophila*. M.J. performed bioinformatic analyses. K.D., E.C.B, B.L, and S.S.G. analyzed the data. M.G. and S.S.G. obtained funding. S.S.G. directed the study and wrote the manuscript. All authors helped edit the final manuscript.

## Figure Legends

**Supplemental Table 1**

Processed RNA-seq data, including lists of up- and down-regulated genes.

## References

Ables, E. T., K. M. Laws and D. Drummond-Barbosa, 2012 Control of adult stem cells in vivo by a dynamic physiological environment: diet-dependent systemic factors in Drosophila and beyond. Wiley Interdiscip Rev Dev Biol 1: 657–674.

Alic, N., T. D. Andrews, M. E. Giannakou, I. Papatheodorou, C. Slack et al., 2011 Genome-wide dFOXO targets and topology of the transcriptomic response to stress and insulin signalling. Mol Syst Biol 7: 502.

Alic, N., M. E. Giannakou, I. Papatheodorou, M. P. Hoddinott, T. D. Andrews et al., 2014a Interplay of dFOXO and two ETS-family transcription factors determines lifespan in Drosophila melanogaster. PLoS Genet 10: e1004619.

Alic, N., J. M. Tullet, T. Niccoli, S. Broughton, M. P. Hoddinott et al., 2014b Cell-nonautonomous effects of dFOXO/DAF-16 in aging. Cell Rep 6: 608–616.

Anderson, L. L., X. Mao, B. A. Scott and C. M. Crowder, 2009 Survival from hypoxia in C. elegans by inactivation of aminoacyl-tRNA synthetases. Science 323: 630–633.

Bandarra, D., J. Biddlestone, S. Mudie, H. A. Muller and S. Rocha, 2014 Hypoxia activates IKK-NF-kappaB and the immune response in Drosophila melanogaster. Biosci Rep 34.

Barretto, E. C., D. M. Polan, A. N. Beevor-Potts, B. Lee and S. S. Grewal, 2020 Tolerance to Hypoxia Is Promoted by FOXO Regulation of the Innate Immunity Transcription Factor NF-kappaB/Relish in Drosophila. Genetics.

Bickler, P. E., and L. T. Buck, 2007 Hypoxia tolerance in reptiles, amphibians, and fishes: life with variable oxygen availability. Annu Rev Physiol 69: 145–170.

Birnbaum, A., X. Wu, M. Tatar, N. Liu and H. Bai, 2019 Age-Dependent Changes in Transcription Factor FOXO Targeting in Female Drosophila. Front Genet 10: 312.

Blaquiere, J. A., W. Lee and E. M. Verheyen, 2014 Hipk promotes photoreceptor differentiation through the repression of Twin of eyeless and Eyeless expression. Dev Biol 390: 14–25.

Borch Jensen, M., Y. Qi, R. Riley, L. Rabkina and H. Jasper, 2017 PGAM5 promotes lasting FoxO activation after developmental mitochondrial stress and extends lifespan in Drosophila. Elife 6.

Bray, N. L., H. Pimentel, P. Melsted and L. Pachter, 2016 Near-optimal probabilistic RNA-seq quantification. Nat Biotechnol 34: 525–527.

Buttgereit, F., and M. D. Brand, 1995 A hierarchy of ATP-consuming processes in mammalian cells. Biochem J 312 (Pt 1): 163–167.

Callier, V., S. C. Hand, J. B. Campbell, T. Biddulph and J. F. Harrison, 2015 Developmental changes in hypoxic exposure and responses to anoxia in Drosophila melanogaster. J Exp Biol 218: 2927–2934.

Centanin, L., A. Dekanty, N. Romero, M. Irisarri, T. A. Gorr et al., 2008 Cell autonomy of HIF effects in Drosophila: tracheal cells sense hypoxia and induce terminal branch sprouting. Dev Cell 14: 547–558.

Centanin, L., P. J. Ratcliffe and P. Wappner, 2005 Reversion of lethality and growth defects in Fatiga oxygen-sensor mutant flies by loss of hypoxia-inducible factor-alpha/Sima. EMBO Rep 6: 1070–1075.

Chen, J., and E. M. Verheyen, 2012 Homeodomain-interacting protein kinase regulates Yorkie activity to promote tissue growth. Curr Biol 22: 1582–1586.

Clegg, J., 1997 Embryos of Artemia franciscana survive four years of continuous anoxia: the case for complete metabolic rate depression. J Exp Biol 200: 467–475.

Danovaro, R., A. Dell’Anno, A. Pusceddu, C. Gambi, I. Heiner et al., 2010 The first metazoa living in permanently anoxic conditions. BMC Biol 8: 30.

Demontis, F., and N. Perrimon, 2010 FOXO/4E-BP signaling in Drosophila muscles regulates organism-wide proteostasis during aging. Cell 143: 813–825.

Dionne, M. S., L. N. Pham, M. Shirasu-Hiza and D. S. Schneider, 2006 Akt and FOXO dysregulation contribute to infection-induced wasting in Drosophila. Curr Biol 16: 1977–1985.

Dobson, A. J., M. Ezcurra, C. E. Flanagan, A. C. Summerfield, M. D. W. Piper et al., 2017 Nutritional Programming of Lifespan by FOXO Inhibition on Sugar-Rich Diets. Cell Rep 18: 299–306.

Doering, K. R. S., X. Cheng, L. Milburn, R. Ratnappan, A. Ghazi et al., 2022 Nuclear hormone receptor NHR-49 acts in parallel with HIF-1 to promote hypoxia adaptation in Caenorhabditis elegans. Elife 11.

Drummond-Barbosa, D., and A. C. Spradling, 2001 Stem cells and their progeny respond to nutritional changes during Drosophila oogenesis. Dev Biol 231: 265–278.

Foe, V. E., and B. M. Alberts, 1985 Reversible chromosome condensation induced in Drosophila embryos by anoxia: visualization of interphase nuclear organization. J Cell Biol 100: 1623–1636.

Fukunaga, K., and N. Shioda, 2009 Pathophysiological relevance of forkhead transcription factors in brain ischemia. Adv Exp Med Biol 665: 130–142.

Giannakou, M. E., M. Goss, M. A. Junger, E. Hafen, S. J. Leevers et al., 2004 Long-lived Drosophila with overexpressed dFOXO in adult fat body. Science 305: 361.

Harrison, J. F., K. J. Greenlee and W. Verberk, 2018 Functional Hypoxia in Insects: Definition, Assessment, and Consequences for Physiology, Ecology, and Evolution. Annu Rev Entomol 63: 303–325.

Harrison, J. F., and G. G. Haddad, 2011 Effects of oxygen on growth and size: synthesis of molecular, organismal, and evolutionary studies with Drosophila melanogaster. Annu Rev Physiol 73: 95–113.

Hemphill, C., E. Pylarinou-Sinclair, O. Itani, B. Scott, C. M. Crowder et al., 2022 Daf-16 mediated repression of cytosolic ribosomal protein genes facilitates a hypoxia sensitive to hypoxia resistant transformation in long-lived germline mutants. PLoS Genet 18: e1009672.

Hochachka, P. W., L. T. Buck, C. J. Doll and S. C. Land, 1996 Unifying theory of hypoxia tolerance: molecular/metabolic defense and rescue mechanisms for surviving oxygen lack. Proc Natl Acad Sci U S A 93: 9493–9498.

Hofmann, G. E., and S. C. Hand, 1994 Global arrest of translation during invertebrate quiescence. Proc Natl Acad Sci U S A 91: 8492–8496.

Holdsworth, M. J., and D. J. Gibbs, 2020 Comparative Biology of Oxygen Sensing in Plants and Animals. Curr Biol 30: R362–R369.

Huang, H., G. Du, H. Chen, X. Liang, C. Li et al., 2011 Drosophila Smt3 negatively regulates JNK signaling through sequestering Hipk in the nucleus. Development 138: 2477–2485.

Hwangbo, D. S., B. Gersham, M. P. Tu, M. Palmer and M. Tatar, 2004 Drosophila dFOXO controls lifespan and regulates insulin signalling in brain and fat body. Nature 429: 562–566.

Junger, M. A., F. Rintelen, H. Stocker, J. D. Wasserman, M. Vegh et al., 2003 The Drosophila forkhead transcription factor FOXO mediates the reduction in cell number associated with reduced insulin signaling. J Biol 2: 20.

Karpac, J., J. Hull-Thompson, M. Falleur and H. Jasper, 2009 JNK signaling in insulin-producing cells is required for adaptive responses to stress in Drosophila. Aging Cell 8: 288–295.

Karpac, J., A. Younger and H. Jasper, 2011 Dynamic coordination of innate immune signaling and insulin signaling regulates systemic responses to localized DNA damage. Dev Cell 20: 841–854.

Kinsey, S. D., J. P. Vinluan, G. A. Shipman and E. M. Verheyen, 2021 Expression of human HIPKs in Drosophila demonstrates their shared and unique functions in a developmental model. G3 (Bethesda) 11.

Kramer, J. M., J. D. Slade and B. E. Staveley, 2008 foxo is required for resistance to amino acid starvation in Drosophila. Genome 51: 668–672.

LaFever, L., A. Feoktistov, H. J. Hsu and D. Drummond-Barbosa, 2010 Specific roles of Target of rapamycin in the control of stem cells and their progeny in the Drosophila ovary. Development 137: 2117–2126.

Lavista-Llanos, S., L. Centanin, M. Irisarri, D. M. Russo, J. M. Gleadle et al., 2002 Control of the hypoxic response in Drosophila melanogaster by the basic helix-loop-helix PAS protein similar. Mol Cell Biol 22: 6842–6853.

Lee, B., E. C. Barretto and S. S. Grewal, 2019 TORC1 modulation in adipose tissue is required for organismal adaptation to hypoxia in Drosophila. Nat Commun 10: 1878.

Lee, W., B. C. Andrews, M. Faust, U. Walldorf and E. M. Verheyen, 2009a Hipk is an essential protein that promotes Notch signal transduction in the Drosophila eye by inhibition of the global co-repressor Groucho. Dev Biol 325: 263–272.

Lee, W., S. Swarup, J. Chen, T. Ishitani and E. M. Verheyen, 2009b Homeodomain-interacting protein kinases (Hipks) promote Wnt/Wg signaling through stabilization of beta-catenin/Arm and stimulation of target gene expression. Development 136: 241–251.

Li, Y., D. Padmanabha, L. B. Gentile, C. I. Dumur, R. B. Beckstead et al., 2013 HIF-and non-HIF-regulated hypoxic responses require the estrogen-related receptor in Drosophila melanogaster. PLoS Genet 9: e1003230.

Liu, G., J. Roy and E. A. Johnson, 2006a Identification and function of hypoxia-response genes in Drosophila melanogaster. Physiol Genomics 25: 134–141.

Liu, L., T. P. Cash, R. G. Jones, B. Keith, C. B. Thompson et al., 2006b Hypoxia-induced energy stress regulates mRNA translation and cell growth. Mol Cell 21: 521–531.

Liu, X., X. Cai, B. Hu, Z. Mei, D. Zhang et al., 2016 Forkhead Transcription Factor 3a (FOXO3a) Modulates Hypoxia Signaling via Up-regulation of the von Hippel-Lindau Gene (VHL). J Biol Chem 291: 25692–25705.

Liu, Y., Y. Wang, X. Li, Y. Jia, J. Wang et al., 2022 FOXO3a in cancer drug resistance. Cancer Lett 540: 215724.

Ma, S., Z. Meng, R. Chen and K. L. Guan, 2019 The Hippo Pathway: Biology and Pathophysiology. Annu Rev Biochem 88: 577–604.

Maiese, K., Z. Z. Chong, Y. C. Shang and J. Hou, 2008 Clever cancer strategies with FoxO transcription factors. Cell Cycle 7: 3829–3839.

Maiese, K., Z. Z. Chong, Y. C. Shang and J. Hou, 2009 FoxO proteins: cunning concepts and considerations for the cardiovascular system. Clin Sci (Lond) 116: 191–203.

Markow, T. A., 2015 The secret lives of Drosophila flies. Elife 4.

Marshall, L., E. J. Rideout and S. S. Grewal, 2012 Nutrient/TOR-dependent regulation of RNA polymerase III controls tissue and organismal growth in Drosophila. EMBO J 31: 1916–1930.

Mendenhall, A. R., B. LaRue and P. A. Padilla, 2006 Glyceraldehyde-3-phosphate dehydrogenase mediates anoxia response and survival in Caenorhabditis elegans. Genetics 174: 1173–1187.

Menuz, V., K. S. Howell, S. Gentina, S. Epstein, I. Riezman et al., 2009 Protection of C. elegans from anoxia by HYL-2 ceramide synthase. Science 324: 381–384.

Morton, D. B., 2011 Behavioral responses to hypoxia and hyperoxia in Drosophila larvae: molecular and neuronal sensors. Fly (Austin) 5: 119–125.

Nakazawa, M. S., B. Keith and M. C. Simon, 2016 Oxygen availability and metabolic adaptations. Nat Rev Cancer 16: 663–673.

Padilla, P. A., and M. L. Ladage, 2012 Suspended animation, diapause and quiescence: arresting the cell cycle in C. elegans. Cell Cycle 11: 1672–1679.

Park, T. J., J. Reznick, B. L. Peterson, G. Blass, D. Omerbasic et al., 2017 Fructose-driven glycolysis supports anoxia resistance in the naked mole-rat. Science 356: 307–311.

Pimentel, H., N. L. Bray, S. Puente, P. Melsted and L. Pachter, 2017 Differential analysis of RNA-seq incorporating quantification uncertainty. Nat Methods 14: 687–690.

Poon, C. L., X. Zhang, J. I. Lin, S. A. Manning and K. F. Harvey, 2012 Homeodomain-interacting protein kinase regulates Hippo pathway-dependent tissue growth. Curr Biol 22: 1587–1594.

Ramirez, J. M., L. P. Folkow and A. S. Blix, 2007 Hypoxia tolerance in mammals and birds: from the wilderness to the clinic. Annu Rev Physiol 69: 113–143.

Raudvere, U., L. Kolberg, I. Kuzmin, T. Arak, P. Adler et al., 2019 g:Profiler: a web server for functional enrichment analysis and conversions of gene lists (2019 update). Nucleic Acids Res 47: W191–W198.

Rideout, E. J., L. Marshall and S. S. Grewal, 2012 Drosophila RNA polymerase III repressor Maf1 controls body size and developmental timing by modulating tRNAiMet synthesis and systemic insulin signaling. Proc Natl Acad Sci U S A 109: 1139–1144.

Romero, N. M., A. Dekanty and P. Wappner, 2007 Cellular and developmental adaptations to hypoxia: a Drosophila perspective. Methods Enzymol 435: 123–144.

Samanta, D., N. R. Prabhakar and G. L. Semenza, 2017 Systems biology of oxygen homeostasis. Wiley Interdiscip Rev Syst Biol Med 9.

Schito, L., and S. Rey, 2018 Cell-Autonomous Metabolic Reprogramming in Hypoxia. Trends Cell Biol 28: 128–142.

Schwenke, R. A., B. P. Lazzaro and M. F. Wolfner, 2016 Reproduction-Immunity Trade-Offs in Insects. Annu Rev Entomol 61: 239–256.

Scott, B., C. L. Sun, X. Mao, C. Yu, B. P. Vohra et al., 2013 Role of oxygen consumption in hypoxia protection by translation factor depletion. J Exp Biol 216: 2283–2292.

Scott, B. A., M. S. Avidan and C. M. Crowder, 2002 Regulation of hypoxic death in C. elegans by the insulin/IGF receptor homolog DAF-2. Science 296: 2388–2391.

Semenza, G. L., 2011 Oxygen sensing, homeostasis, and disease. N Engl J Med 365: 537–547.

Semenza, G. L., 2014a Hypoxia-inducible factor 1 and cardiovascular disease. Annu Rev Physiol 76: 39–56.

Semenza, G. L., 2014b Oxygen sensing, hypoxia-inducible factors, and disease pathophysiology. Annu Rev Pathol 9: 47–71.

Short, S. M., M. F. Wolfner and B. P. Lazzaro, 2012 Female Drosophila melanogaster suffer reduced defense against infection due to seminal fluid components. J Insect Physiol 58: 1192–1201.

Sivaraj, K. K., B. Dharmalingam, V. Mohanakrishnan, H. W. Jeong, K. Kato et al., 2020 YAP1 and TAZ negatively control bone angiogenesis by limiting hypoxia-inducible factor signaling in endothelial cells. Elife 9.

Slack, C., M. E. Giannakou, A. Foley, M. Goss and L. Partridge, 2011 dFOXO-independent effects of reduced insulin-like signaling in Drosophila. Aging Cell 10: 735–748.

Sriskanthadevan-Pirahas, S., R. Deshpande, B. Lee and S. S. Grewal, 2018 Ras/ERK-signalling promotes tRNA synthesis and growth via the RNA polymerase III repressor Maf1 in Drosophila. PLoS Genet 14: e1007202.

Steinmetz, E. L., D. N. Dewald and U. Walldorf, 2021 Drosophila Homeodomain-Interacting Protein Kinase (Hipk) Phosphorylates the Hippo/Warts Signalling Effector Yorkie. Int J Mol Sci 22.

Supek, F., M. Bosnjak, N. Skunca and T. Smuc, 2011 REVIGO summarizes and visualizes long lists of gene ontology terms. PLoS One 6: e21800.

Teleman, A. A., V. Hietakangas, A. C. Sayadian and S. M. Cohen, 2008 Nutritional control of protein biosynthetic capacity by insulin via Myc in Drosophila. Cell Metab 7: 21–32.

Teodoro, R. O., and P. H. O’Farrell, 2003 Nitric oxide-induced suspended animation promotes survival during hypoxia. EMBO J 22: 580–587.

Tettweiler, G., J. A. Blaquiere, N. B. Wray and E. M. Verheyen, 2019 Hipk is required for JAK/STAT activity during development and tumorigenesis. PLoS One 14: e0226856.

Tettweiler, G., M. Miron, M. Jenkins, N. Sonenberg and P. F. Lasko, 2005 Starvation and oxidative stress resistance in Drosophila are mediated through the eIF4E-binding protein, d4E-BP. Genes Dev 19: 1840–1843.

Texada, M. J., A. F. Jorgensen, C. F. Christensen, T. Koyama, A. Malita et al., 2019 A fat-tissue sensor couples growth to oxygen availability by remotely controlling insulin secretion. Nat Commun 10: 1955.

van den Beucken, T., M. Koritzinsky and B. G. Wouters, 2006 Translational control of gene expression during hypoxia. Cancer Biol Ther 5: 749–755.

Verheyen, E. M., S. Swarup and W. Lee, 2012 Hipk proteins dually regulate Wnt/Wingless signal transduction. Fly (Austin) 6: 126–131.

Webb, A. E., and A. Brunet, 2014 FOXO transcription factors: key regulators of cellular quality control. Trends Biochem Sci 39: 159–169.

Wingrove, J. A., and P. H. O’Farrell, 1999 Nitric oxide contributes to behavioral, cellular, and developmental responses to low oxygen in Drosophila. Cell 98: 105–114.

Wong, D. M., Z. Shen, K. E. Owyang and J. A. Martinez-Agosto, 2014 Insulin-and warts-dependent regulation of tracheal plasticity modulates systemic larval growth during hypoxia in Drosophila melanogaster. PLoS One 9: e115297.

Wong, K. K. L., J. Z. Liao and E. M. Verheyen, 2019 A positive feedback loop between Myc and aerobic glycolysis sustains tumor growth in a Drosophila tumor model. Elife 8.

Wu, Z., and K. L. Guan, 2021 Hippo Signaling in Embryogenesis and Development. Trends Biochem Sci 46: 51–63.

Zhou, D., J. Xue, J. C. Lai, N. J. Schork, K. P. White et al., 2008 Mechanisms underlying hypoxia tolerance in Drosophila melanogaster: hairy as a metabolic switch. PLoS Genet 4: e1000221.

